# An Updated Investigation Prior To COVID-19 Vaccination Program In Indonesia: Full-Length Genome Mutation Analysis Of SARS-CoV-2

**DOI:** 10.1101/2021.01.26.426655

**Authors:** Reviany V. Nidom, Setyarina Indrasari, Irine Normalina, Astria N. Nidom, Balqis Afifah, Lestari Dewi, Andra Kusuma Putra, Arif N. M. Ansori, Muhammad K. J. Kusala, Mohammad Y. Alamudi, Chairul A. Nidom

## Abstract

**Introduction:** Indonesia kick-started the big project of COVID-19 vaccination program in January 2021 by employed vaccine to the president of Indonesia. The outbreak and rapid transmission of COVID-19 have endangered the global health and economy. This study aimed to investigate the full-length genome mutation analysis of 166 Indonesian SARS-CoV-2 isolates as 12 January 2021.

**Methods:** All data of isolates was extracted from the Global Initiative on Sharing All Influenza Data (GISAID) EpiCoV database. CoVsurver was employed to investigate the full-length genome mutation analysis of all isolates. Furthermore, this study also focused on the unlocking of mutation in Indonesian SARS-CoV-2 isolates S protein. WIV04 isolate that was originated from Wuhan, China was used as a virus reference according to CoVsurver default. All data was visualized using GraphPad Prism software, PyMOL, and BioRender.

**Results:** This study result showed that a full-length genome mutation analysis of 166 Indonesian SARS-CoV-2 isolates was successfully discovered. Every single mutation in S protein was described and then visualised by employing BioRender. Furthermore, it also found that D614G mutation appeared in 103 Indonesian SARS-CoV-2 isolates.

**Conclusion:** To sum up, this study helps to observe the spread of the COVID-19 transmission. However, it would like to propose that the epidemiological surveillance and genomics studies might be improved on COVID-19 pandemic in Indonesia.

## INTRODUCTION

SARS-CoV-2 firstly occurred in China and then transmitted sporadically worldwide. In March 2020, WHO announced that the infection was a pandemic. COVID-19 outbreak and rapid transmission have endangered global health and economy^1^. This crisis has called for an extensive scientific mobilization of studies on SARS-CoV-2 focusing its clinical aspects, characteristics, and its mechanism of transmission, with the ultimate aim of counteracting the devastating outcomes^2,3^. Recently, there have been around 92 million people globally who have been infected by the seventh coronavirus and more than 2 million deaths as the fast result of this pandemic. Specifically, there are more than 850,000 cases and around 25,000 people died in Indonesia. These data are derived from Johns Hopkins University online website that tracks COVID-19 cases in real-time^4^.

Meanwhile, on coronaviruses themselves, the family *Coronaviridae* is categorised into four different genera: *Gammacoronavirus, Deltacoronavirus, Betacoronavirus*, and *Alphacoronavirus*. Both animals and humans can be infected by coronaviruses^5^. Moreover, SARS-CoV-2 genome is a single-stranded positive-sense RNA of roughly 30,000 nucleotides. There are four structural proteins encoded by the genome and spike (S) protein is the most important one^6^. It is because S protein is the primary target antigen in the SARS-CoV-2 vaccine^7^. Previously, the candidate for a peptide-based vaccine against the virus based on the four structural proteins was identified^8,9^. In addition, the interaction between the host and the virus that causes infection involves a complex response of immune system^10,11^. On the other hand, we demonstrated the paradoxical phenomenon called antibody-dependent enhancement (ADE) within the Indonesian isolates^12^. Therefore, ADE has become a tipping point in the cultivation of antibody-based therapies and vaccines^10,11^.

Recently, scientists are attempting to generate vaccines to fight against SARS-CoV-2 worldwide, with protein-based vaccines as the most advanced types and the private sector is at the forefront of this study^13,14,15,16^. Unexpectedly/uncontrollably, the mutation rate of *Coronaviridae* is reminding high. Various studies reported the mutation might be implicated the efficacy of vaccines or other therapeutic strategies^17,18,19^. Furthermore, the newest type of SARS-CoV-2 lineage occurred in UK at the end of 2020 and this case predicted as sporadically would increase the number of the COVID-19 patients in UK. Thus, investigating the full-length genome mutation analysis of 166 Indonesian SARS-CoV-2 isolates became the goal of this study.

## MATERIALS AND METHODS

### SARS-CoV-2 Isolates

The data extraction of all Indonesian isolates (166 isolates) with SARS-CoV-2 from the Global Initiative on Sharing All Influenza Data (GISAID) EpiCoV database was completed on 12 January 2021. This study only used the complete genome and high coverage criteria according to the GISAID EpiCoV standard. All of 166 Indonesian SARS-CoV-2 isolates were derived from various provinces in Indonesia, for example: Special Region of Aceh, North Sumatra, Lampung, Banten, Special Capital Region of Jakarta, West Java, Central Java, Special Region of Yogyakarta, East Java, and so on. This study identified the total isolates in every province, GISAID clades, and lineages. All data were visualized using GraphPad Prism software. Furthermore, WIV04 isolate (GISAID clade: L; lineage: B) collected from a female retailer at Huanan Seafood Wholesale Market and submitted by the Wuhan Institute of Virology, Chinese Academy of Sciences, China was applied as a virus reference based on CoVsurver default.

### Full-Length Genome Mutation Analysis

Every single gene of 166 Indonesian SARS-CoV-2 isolates was investigated, such as NSP1-16, S, NS3, E, M, NS6, NS7a, NS7b, NS8, and N. Also, D614G mutation and S protein mutations in all isolates were identified. We employed CoVsurver to investigate the full-length genome mutation analysis of all isolates. All data were visualized using GraphPad Prism software and BioRender.

### 3D Structure Visualization

In this study, we rendered 3D structure visualization of SARS-CoV-2 S protein by applying SWISS-MODEL web server and PyMOL v2.4. Then, the schematic diagram was edited with BioRender. This method was functioned to easily identify the location of various mutations in SARS-CoV-2 S protein.

## RESULTS AND DISCUSSION

Until at the end of 2019, there were six identified coronaviruses to be causative agents of infection in humans. The seventh, SARS-CoV-2, emerged in China^1^. To date, according to the CSSE at Johns Hopkins University, there are approximately 95 million people infected with the virus globally^4^. Moreover, the reports said that human-to-human transmission has occurred and WHO has acknowledged the chance of aerosol infection^20^. Based on the data, the novel virus has been collected from saliva, throat, bronchoalveolar-lavage, oropharyngeal swab, nasopharyngeal swab, and sputum.

The data of 166 Indonesian SARS-CoV-2 isolates were retrieved from the database prior to the Indonesia’s COVID-19 vaccination program kick-started on 13 January 2021. The database was the collective results from nasopharyngeal swab, oropharyngeal swab and nasopharyngeal swab, and sputum methods which were used to collect the virus samples and submitted by many collaborations among research centres and universities in Indonesia. The 166 isolates were originated from various provinces in Indonesia, such as Special Region of Aceh, North Sumatra, Riau Islands, Lampung, Banten, Special Capital Region of Jakarta, West Java, Central Java, Special Region of Yogyakarta, East Java, South Kalimantan, North Kalimantan, Central Kalimantan, East Kalimantan, North Sulawesi, Bali, West Nusa Tenggara, East Nusa Tenggara, North Papua, West Papua, dan Papua (Figure 1).

**Figure 1.**
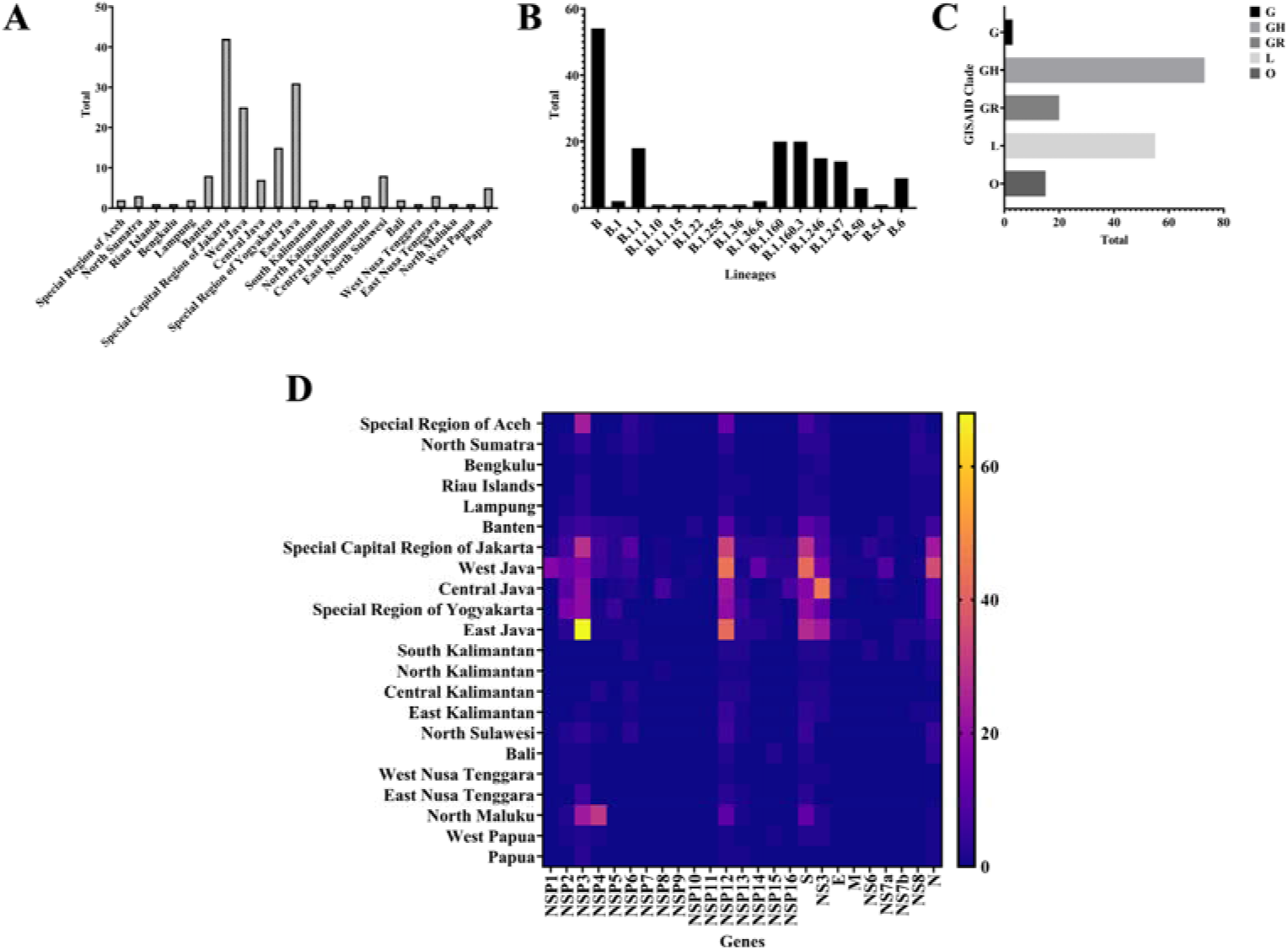
All Indonesian SARS-CoV-2 isolates data. A. The origin of isolates; B. The lineages distribution of the isolates; C. The GISAID EpiCoV clades; and D. The full-length genome mutation analysis of all isolates used in this study. All data were visualized using GraphPad Prism software.

Recent updates show that the GISAID EpiCoV database has acknowledged seven subtypes of SARS-CoV-2, specifically V, S, O, L, GR, GH, GV, and G clades. Significantly, the isolates from Indonesia in this study were grouped in the G, GH, GR, L, and O clades. Similarly, this study also reveals those above lineages of SARS-CoV-2 isolates in Indonesia according to the accumulated data from GISAID EpiCoV (Figure 1). In this study, we revealed 14 lineages from 166 isolates (Figure 1). However, based on this data, there are no novel lineages identified related to new variants. In contrast, many reports stated that novel variants of SARS-CoV-2 occurred in various countries, such as UK, Brazil and South Africa^21,22,23.24,25^. Consequently, these novel variants might be more transmissible and suspected to be accountable for the rise of COVID-19 patient numbers in those countries^23,24^. On the other hand, the previous studies about the molecular phylogenetic tree revealed that the relationship of SARS-CoV-2 and other *Coronaviridae* is based on the four structural protein genes. In accordance with this, SARS-CoV-2 is considered to be the closest to *Rhinolophus affinis* coronavirus RaTG13 and followed by pangolin coronavirus^26^. Thus, Malayan pangolin is assumed as the intermediate host before infecting to humans^27^.

Meanwhile, S protein mediates the entry and the new virus membrane fusion and is the main target for many studies of antiviral drugs and vaccines^28,29^. S1 and S2 are the two domains of the virus S protein (Figure 2). S1 is conscientious for binding to host cellular receptors. Besides the efficacy of several therapies which include disrupting protease inhibitors, small RNAs, neutralizing antibodies, fusion blockers, S protein inhibitors, ACE2 blockers, however, the *in vitro* studies on S protein inhibitors have been unsatisfactory^30^. Many methods have been employed to produce vaccines using S protein as an antigen^8,31^. Thus, it is very important to investigate the S protein from Indonesian SARS-CoV-2 isolates.

**Figure 2.**
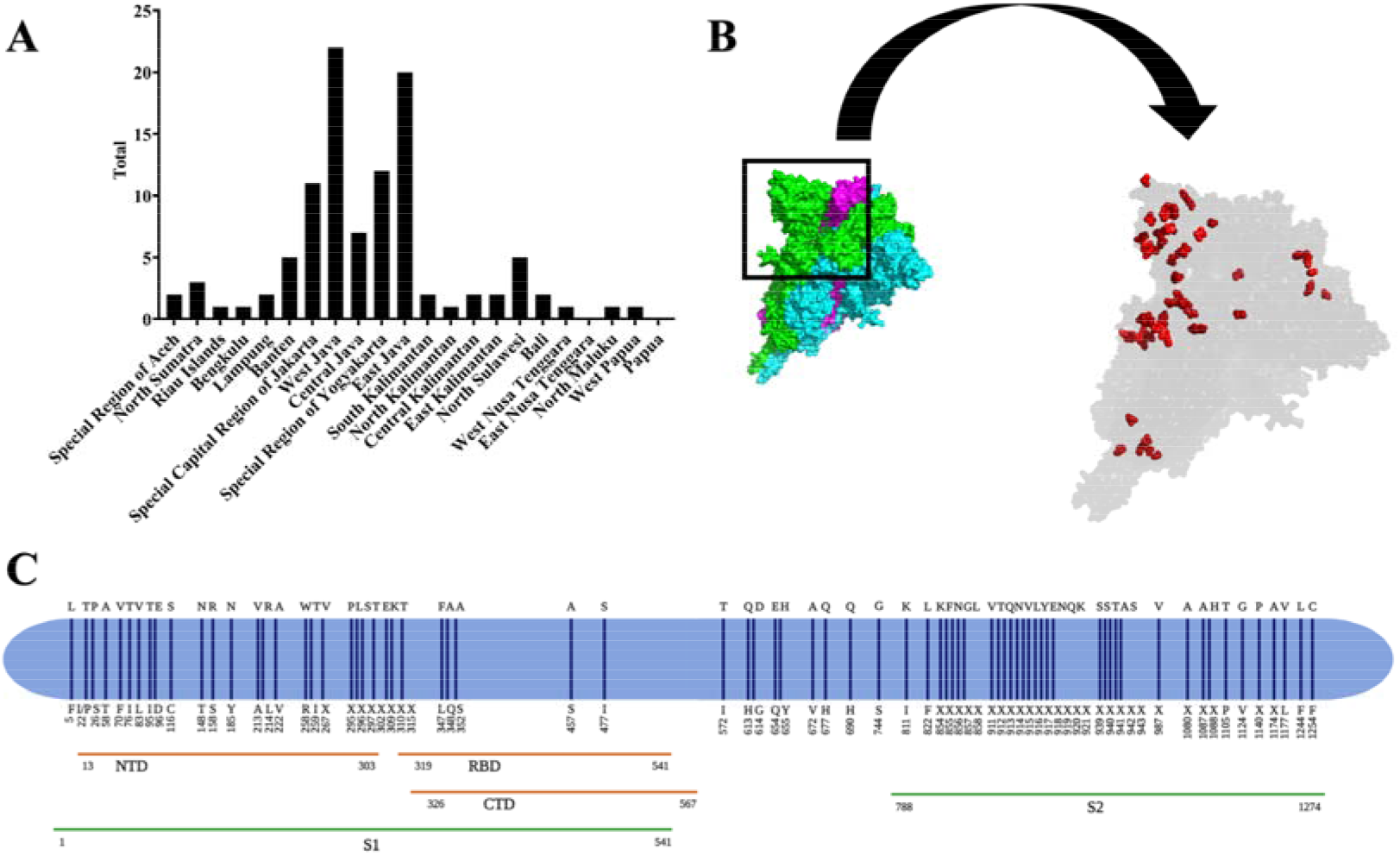
Mutation analysis originated from S protein of Indonesian SARS-CoV-2. A. Distribution of D614G mutation from various provinces in Indonesia; B. 3D structure visualization of SARS-CoV-2 S protein. Red dots are the location of various mutations in S protein; and C. The mapping of amino acid mutation sites in S protein of 166 Indonesian SARS-CoV-2 isolates.

Scientists have demonstrated that mutations occur in the virus genome globally^7,9,32,33^. Previously, Phan *et al*. performed a genetic analysis in 86 virus genomes and reported many mutations. One of the most important mechanisms proposed for the evolution of viruses in nature is nucleotide substitution^7^. Meanwhile, Yadav *et al*. also reported a study to analyse the first two virus isolates from India^34^. In line with this, Garcés-Ayala *et al*. conducted a study with the reference sequence for fully describing the novel SARS-CoV-2 complete genome in Mexico^35^. Concurrently, Khailany *et al*. retrieved 94 SARS-CoV-2 genomes and checked the molecular variation between them^36^. Whilst, Kim *et al*. also revealed that the quick transmission and infectivity of the virus correlated with specific mutations in the genome^37^. Simultaneously, this study reported various S protein mutations such as A222, S477, D614, Q677, and so on. Further research was highly considered that S protein mutations to affect vaccination program worldwide^38,39,40^.

Recent publications shows that one of the most notable amino acid mutations is D614G^12,13^. Based on these recent studies, the virus virulence and the increase of viral loads in COVID-19 patients characterise the occurrence of D614G mutation^40,41^. While, based on the current available information, the infectivity as well as the receptor binding, fusion activation, or ADE enhancement can be influenced by D614G mutation in several ways^10,11,12^. An antibody escape is considered as another mutation mechanism like the upcoming form of D614G which can be accelerated by an antigenic drift. If the sensitivity of neutralizing antibody can be affected by D614G mutation in SARS-CoV-2 or vice versa, then, the ADE activity also can be monitored in the SARS-CoV study, thus, D614G can be considered as an intermediate antibody escape which puts people to be more vulnerable for second infections^12,40,42^.

In this study, D614G mutation was detected in 103 Indonesian SARS-CoV-2 isolates. All the isolates were mostly from the West Java, East Java, Special Region of Yogyakarta, Special Capital Region of Jakarta, and Central Java, respectively (Figure 2). This result is in line with a study by Zhang *et al*. on D614G mutation which discovered that S1 residue 614 is in a close proximity to S2 domain. An altered release or shedding of S1 domain after cleavage at S1/S2 junction might be displayed by the ratio between S1 and S2 domains in the virion. Glycine Amino Acid found at residue 614 of S protein G614 secures the interaction between S1 and S2 domains and limits S1 shedding. D614G mutation has been previously speculated in raising an open configuration of S protein that is more advantageous to ACE2 association^40^. Therefore, SARS-CoV-2 S protein D614G mutation is highly believed in promoting the virion spike density and infectivity and it is also highly speculated that this mutation might be influence further mutations. In addition, the previous study also reported that the type of mutation emerged in the virus isolates were originated from canine, environment, *Felis catus, Mus musculus, Mustela lutreola*, and *Panthera tigris jacksoni*^12^.

In the meantime, compared to most other microorganisms, the rates of RNA viruses’ mutations are much higher^43^. An elevated mutation rate can lead to an increase in virulence and a high potential for adaptive evolution^43,44^. This capability boosts the chance of zoonotic viral pathogens to establish human-to-human transmission and permits them to enhance their virulence^45^.

This study provides the fundamental data for accomplished studies into the medication and prevention of COVID-19. Indonesian SARS-CoV-2 genomic data extraction would be valuable in vaccine construction and options in medication. In fact, mining the data of the Indonesian SARS-CoV-2 variants and molecular epidemiology could enable the mapping of its origin and the tracking of its transmission^46^. In line with this, the sequence investigation performs an important role in viral surveillance, public health policy problems, and host identification^46,47^. Thus, high-speed detection of mutations from the Indonesian SARS-CoV-2 is mandatory in the unlocking to the COVID-19 pandemic in Indonesia.

Currently, the availability of COVID-19 vaccine is limited, not many people can access it. In countries that have not implemented large-scale active case testing and isolation, controlling the spread of the virus can be a challenge. In this case, transmission suppression relies primarily on the community adherence to non-pharmacological strategies such as social distancing, the mandatory mask using, and hand washing^48,49,50,51^. Inevitably, as the result of SARS-CoV-2 outbreaks, many countries declared medical emergency which led to economic emergency since those countries enforcing limited or strict mobility both regionally and nationally^52,53,54^. Thus, in regulating and containing further transmission of COVID-19, it is a fundamental move to discover the characteristics of SARS-CoV-2 genome and constitute the systems for observing SARS-CoV-2 during this pandemic. Furthermore, the recognition of genotypes related to temporal infectious clusters and specific geographic areas recommends that the employment of genomic data is highly recommended in observing and tracking the further spreading of SARS-CoV-2. Likewise, researchers might be able to introduce the origin of a specific variant and observe the virus transmission by acknowledging the specific SARS-CoV-2 variants and connecting them using a molecular epidemiology approach. Ergo, it can be argued that this study might become an important tool in regulating the COVID-19 pandemic in Indonesia.

## CONCLUSION

In conclusion, this study successfully identified the full-length genome mutation analysis of 166 Indonesian SARS-CoV-2 isolates. This study helps in observing the spread of the COVID-19 transmission. However, we proposed that the epidemiological surveillance and genomics studies might be improved on COVID-19 pandemic in Indonesia.

## ACKNOWLEDGMENT

This study was fully supported by the Professor Nidom Foundation, Indonesia (Grant Number: 005/PNF-RF/07/2020). We would like to take this opportunity to express our deepest condolences to the victims of COVID-19. We would also like to pay tribute to our frontline heroes who have fought continuously to this day. Thankfully, we also recognize the authors from the originating and submitting laboratories of GISAID database on which the samples were derived from. We would like to thank to National Institute of Health Research and Development, Ministry of Health, Indonesia (Balitbangkes Kemenkes) for the supports. Additionally, we also thank Dewi Sartika, M.Ed. for her help in editing the manuscript.

